# Historic and prehistoric human-driven extinctions have reshaped global mammal diversity patterns

**DOI:** 10.1101/017368

**Authors:** S. Faurby, J-C. Svenning

## Abstract

**Aim:** To assess the extent to which humans have reshaped Earth’s biodiversity, by estimating natural ranges of all late Quaternary mammalian species, and to compare diversity patterns based on these with diversity patterns based on current distributions.

**Location:** Globally

**Methods:** We estimated species, functional and phylogenetic diversity patterns based on natural ranges of all mammalian species (n=5747 species) as they could have been today in the complete absence of human influence through time. Following this we compared macroecological analyses of current and natural diversity patterns to assess if human-induced range changes bias for evolutionary and ecological analyses based on current diversity patterns.

**Results:** We find that current diversity patterns have been drastically modified by humans, mostly due to global extinctions and regional to local extirpations. Current and natural diversities exhibit marked deviations virtually everywhere outside sub-Saharan Africa. These differences are strongest for terrestrial megafauna, but also important for all mammals combined. The human-induced changes led to biases in estimates of environmental diversity drivers, especially for terrestrial megafauna, but also for all mammals combined.

**Main conclusions:** Our results show that fundamental diversity patterns have been reshaped by human-driven extinctions and extirpations, highlighting humans as a major force in the Earth system. We thereby emphasize that estimating natural distributions and diversities is important to improve our understanding of the evolutionary and ecologically drivers of diversity as well as for providing a benchmark for conservation.

## (A). Introduction

Human activities increasingly affect the whole Earth system (Crutzen, 2002), driving an ongoing global mass extinction (Barnosky *et al.*, 2011) and massive global environmental changes (IPCC, 2013), with a looming planetary biosphere state shift on the horizon (Barnosky *et al.*, 2011). A largely overlooked consequence of these anthropogenic global transformations is that they may also influence our ability to understand the factors that have generated and maintained Earth’ s biodiversity, one of the most important questions for contemporary science (Pennisi, 2005). Mammals represent one of the most studied organism groups, and the current diversity and general distribution of most mammal groups is well known (Schipper *et al*., 2008). This knowledge has been used in numerous studies analyzing patterns in species, functional, or phylogenetic diversity (Safi *et al*., 1011; Huang *et al*., 1012; Jetz & Fine, 2012; Mazel *et al*., 2014), as well as conservation studies (Sodhi *et al*., 2010). However, mammals are also one of the organism groups that have been influenced the most by human activities, such as habitat loss and hunting (Schipper *et al*., 2008; Spear & Chown, 2008; Sandom *et al*., 2014). Therefore, it is not clear to what extent the current diversity patterns still reflect natural patterns or are biased by anthropogenic extinctions, extirpations, and introductions. The extent to which our knowledge on the natural drivers of mammal diversity is biased by these human-induced changes is unknown.

There is ample evidence that human activities have strongly the distributions of a number of mammal species during the last few thousand years. These modifications involve range contractions (Short & Smith, 1994; Laliberte & Ripple 2004), but also extinctions, notably on islands (Turvey & Fritz, 2011), and to a lesser extent also old introductions of species that are now thought of as native (Bover & Alcover, 2008). In addition, accumulating evidence indicates that humans have been a major driver of severe Late Pleistocene and early Holocene large-mammal extinctions (Sandom *et al*., 2014). These human-induced changes raise the question of how diversity patterns would look today without any human modifications of species distributions in the Late Pleistocene, early Holocene, or historic time, i.e., given the present-natural ranges *sensu* Peterken (1977) for all late Quaternary mammals. The term “present-natural” refers to the state that a phenomenon would be in today in the complete absence of human influence through time; for simplicity, we hereafter refer to this concept by the term ‘natural’.

The majority of basic ecological and evolutionary studies on diversity patterns in mammals have aimed to test hypotheses for natural diversity patterns, but testing them on current diversity patterns (Sandom *et al.*, 2013). The potential problems that such anthropogenic effects may cause for macro-ecological studies have previously been pointed out (Blackburn & Gaston, 1998). Yet, no previous study has systematically estimated the natural distribution of all species within larger clades, and empirical studies have been forced to ignore this issue. If current diversity patterns have been re-shaped by human activities, this may confound conclusions from studies of natural drivers. Therefore, knowing how different current diversity patterns are from the natural patterns is important, but no study has assessed this. The goal of this paper is to estimate natural diversity patterns of mammals (i.e., as they would have been given the natural distributions of all species) in order determine how much human activities have reshaped contemporary diversity patterns. Understanding how current diversity patterns are shaped by human activities is not just crucial for diversity studies, but also essential for conservation management, notably for providing base-lines for restoration efforts (Donlan *et al.*, 2005).

As argued by Devictor *et al.* (2010), no single diversity measure fully captures all biologically relevant elements of diversity; therefore, we estimated natural patterns not just for taxonomic diversity (species richness), but also for phylogenetic and functional diversity. The use of multiple diversity measures is especially interesting in this context because all previous analyses of pre-historic faunal losses have focused solely on taxonomic richness (Sandom *et al*., 2014; Johnson, 2002). We also analyzed the effects of human-induced changes in diversity on estimates of relationships between diversity and key environmental factors in order to investigate the extent to which analyses based on current diversity patterns may be biased. We expect that the largest differences will be found for larger animals because both the Late Pleistocene extinctions and the more recent range contractions have disproportionally affected larger species (Koch & Barnosky, 2006; Ripple *et al*., 2014). We also expect pronounced geographic variation in the difference between current and natural diversity due to variation in human impact between geographic regions (De Thoisy *et al*., 2010), as well as higher sensitivity of island or island-like faunas (Sandom *et al*., 2014; Duncan *et al*., 2013). In addition, some habitat types may have higher sensitivity than others, with large, dense forests and steep mountainous areas being less vulnerable (Johnson, 2002) and desert regions being more vulnerable (Yeakel, 2014).

## (A) Materials and methods

### (B) Range modifications

The taxonomy for this study is the same as in a recent mammal phylogeny (Faurby & Svenning, 2015). For all species, we attempted to estimate their potential current distributions as they would potentially be today if they had not been modified by humans, i.e., their natural ranges (Peterken, 1977). Climatic data were used to estimate this range for some species, but this is not identical to the climatic potential range because we only aimed to identify the areas the species would have been able to inhabit without human interference, not the entire possible potential range based on the estimated niche. Therefore, natural dispersal constraints, biotic constraints, and non-climatic abiotic limiting factors were also taken into account when estimating natural ranges. We systematically inspected the IUCN ranges (Schipper *et al*., 2008) of all species suspected to have had anthropogenically-induced range changes based on red-list category (red-listed as vulnerable, endangered, critically endangered, extinct, extinct in the wild, or data deficient: 2302 species), body size (all species larger than 1 kg: 627 additional species), or occurrence in large isolated island-like systems (Australia, New Guinea, or Madagascar: 340 additional species). These ranges were modified accordingly when evidence for anthropogenic range changes was found. The remaining species were not systematically investigated, but their ranges were modified whenever we found evidence for human-caused range changes. We modified the ranges of 1085 species, though for 85 of them the range modifications were too small to affect our analyses at our chosen grid size (110 × 110 km cells). A total of 260 species were not covered by IUCN because they went extinct prior to 1500 AD, but were included in our analysis because they have gone extinct within the last 130,000 years. By doing this we implicitly assumed that all global and continental extinctions during this period were caused by humans rather than natural phenomena, such as climatic variations. We acknowledge that this may not be true in all cases, as the probability of extinction prior to human contact seems high for a few species, such as the giant Caribbean rodent *Amblyrhiza inundata* (blunt-toothed giant hutia) (Biknevicus *et al*., 1993). Still, the evidence is overwhelming for strong human involvement in most of these extinctions (Sandom *et al*., 2014; Turvey & Fritz 2011). Our methodology did of course allow for natural regional extinctions, notably due to climate changes, e.g., the disappearance of *Ovibos moschatus* (musk-ox) and *Gulo gulo* (wolverine) and other cold-adapted species from Southern Europe after the end of the ice age (Álvarez-Lao & García, 2010).

Overall, the modifications led to a change from a total of 1,983,482 occurrences to 2,212,446 occurrences in 110 × 110 km cells (1,073,129 to 1,298,365 occurrences for non-marine mammals and 67,306 to 242,960 occurrences for megafauna). A detailed explanation of the different types of modifications can be found in supplementary methods.

### (B) Estimated diversity

All diversities were estimated on a Behrman projection of the world with equally sized grid cells with 360 columns (i.e., 1° by 1° cells at the equator, roughly equal to 110 × 110 km grid cells). Diversities were estimated for five mammalian subgroups: 1) *all species*, 2) *non-marine species*, 3) *terrestrial species*, 4) *large terrestrial species*, and 5) *terrestrial megafauna*. For *non-marine species*, species coded as exclusively marine by IUCN (most whales and two manatees), as well as pinnipeds coded as “marine and terrestrial” and three effectively marine non-pinniped carnivores (*Ursus maritimus* (polar bear), *Enhydra lutris* (sea otter) and *Lontra felina* (marine otter)), were removed, whereas species coded as “freshwater and marine”, such as *Phoca vitulina* (harbor seal) or manatees of the genus *Trichechus*, were deleted from all fully or partially marine cells. For *terrestrial species* all manatees, whales, pinnipeds, and bats were removed from *non-marine species*, and this list was further restricted in *large terrestrial species* to only include species larger than 10 kg, the definition of megafauna used by Sandom *et al*. (2014), and in *terrestrial megafauna* to only include species greater than 44.5 kg, the classical definition of megafauna used by many studies (Barnosky *et al*., 2004).

Analyses were performed on species, phylogenetic, and functional diversity. The phylogenetic diversity of each cell was defined as the median tree length of the species in the cell based on 100 trees from the posterior distribution of the phylogeny (Faurby & Svenning, 2015). Our treatment of functional diversity is a multidimensional version of the bin-filling approach of (Huang *et al*., 2012). For the functional diversity analyses of *all species* and *non-marine species*, we focused on the three dimensions of niche space, habitat, body size, and diet, whereas the analysis of the three terrestrial subsets only focused on body size and diet. Details of the estimation can be found in supplementary methods.

Three diversities were estimated for each cell: 1) current diversity, 2) natural diversity of historically extant species, and 3) total natural diversity. *Current diversity* was defined as the diversity following IUCN, excluding species ranges coded as introduced and species ranges coded as extinct or possibly extinct (*5*). *Natural diversity of historically extant species* was generally the natural diversity of all species accepted by IUCN, meaning that species ranges coded as extinct or possibly extinct by IUCN, as well as our modified ranges of species accepted by IUCN, were included, but species that went globally extinct prior to 1500 AD were not included. In addition, for species that went continentally extinct (with Europa and Asia considered the same continent) prior to 1500 AD, the natural distributions on these continents were removed. Therefore, the natural distribution of *Equus ferus* in North America, South America, and Africa, *Camelus dromedarius* in Africa and Eurasia, *Cuon alpinus* and *Saiga tatarica* in North America, *Bos primigenius* in Africa, and *Crocuta crocuta, Hippopotamus amphibius, Macaca sylvanus,* and *Ovibos moschatus* in Eurasia were removed. *Total natural diversity* included the natural distribution of all species accepted by IUCN, including the ranges on the continents mentioned above, but also included the natural distribution of the 260 pre-historically extinct species. In the main article we focus on the differences between *current diversity* and *total natural diversity*, and we refer to *total natural diversity* simply as *natural diversity* throughout the main article. Separate maps showing the patterns for *Natural diversity of historically extant species* are shown in the supplementary figures.

We also estimated the total deficit (the difference between *current diversity* and *total natural diversity* relative to the *total natural diversity*), the historic loss (the difference between *current diversity* and *natural diversity of historically extant species* relative to the *total natural diversity*), and the pre-historic loss (the difference between the *natural diversity of historically extant species* and the *total natural diversity* relative to the *total natural diversity*). These terms are defined temporally, and the vast majority of pre-historic loss occurred earlier than historic loss, but we note that a limited temporal overlap between the two exists. For example, the massive pre-historic loss in the Caribbean mainly occurred within the middle to late Holocene (Steadman *et al*., 2005), whereas most of the range contractions of *Panthera pardus* in Europe occurred earlier (Sommer & Benecke, 2006). In addition, for some species the loss in range was a slow and gradual process, perhaps best exemplified by *Equus ferus*. The decline in this species started near the end of the last ice age with continental extinctions in North and South America (Sandom *et al.*, 2014), whereas the last wild specimen in Europe died in the 19^th^ century(Nowak, 1999) and the last wild specimen of the species globally died in the 20^th^ century (Schipper *et al*., 2008).

### (B) Statistical analysis of diversity

We analyzed the geographic variation in diversity for a total of 45 different analyses: all combinations of the three diversities (species, phylogenetic, and functional), five datasets (terrestrial megafauna, large terrestrial mammals, all terrestrial mammals, non-marine mammals, and all mammals) and current, IUCN natural, and total natural diversity. Our analysis had nine variables comprising seven main effects and two interactions. The effects were: 1) elevation range, 2) annual temperature (Hijmans *et al*., 2005), 3) logarithm transformed annual precipitation (Hijmans *et al*., 2005), 4) precipitation seasonality (Hijmans *et al*., 2005), 5) temperature seasonality (Hijmans *et al*., 2005), 6) NDVI (Tucker *et al*., 2005), and 7) “open areas”. The last was a dummy variable separating all non-forest cells (defined as areas with Tropical and Subtropical Grasslands, Savannas and Shrublands, Temperate Grasslands, Savannas and Shrublands, Flooded Grasslands and Savannas, Montane Grasslands and Shrublands, Tundra, or Deserts and Xeric Shrublands (Olson, 2001)) from forest cells (defined as Tropical and Subtropical Moist Broadleaf Forests, Tropical and Subtropical Dry Broadleaf Forests, Tropical and Subtropical Coniferous Forests, Temperate Broadleaf and Mixed Forests, Temperate Coniferous Forest, Boreal Forests/Taiga, Mediterranean Forests, Woodlands and Scrubs, or Mangroves (Olson, 2001)). The interactions were: 8) annual temperature and open areas and 9) annual temperature and annual precipitation. The interactions were intended to take into account that diversity may be different in forest and non-forest biomes, especially for megafauna because such species may not have access to the plant resources in the canopy. All parameters except “open areas” were standardized to have a mean of 0 and a standard deviation of 1.

In order to remove spatial autocorrelation in the data, we analyzed the data based on the SARerr model, which has been suggested to be a suitable method of minimizing autocorrelation (Kissling & Carl, 2008). SAR models are computing intensive; therefore, for computational reasons, we chose to perform them using cells equivalent to 4° squares at the equator. We tried neighborhoods for each SAR model of between 1 and 8 neighbors and selected the best model based on AIC (the chosen models had between 4 and 6 neighbors). Next, we estimated the overall model performance by calculating the square of the correlation between the predicted (only the predictor, not the spatial parts) and raw values. We refer to this as pseudo-R^2^ throughout the paper even though several different estimates of model fit are frequently referred to as pseudo-R^2^ (UCLA: Statistical Consulting Group, 2014). All p-values were calculated by Wald’ s tests. In order to make comparisons between different models easier, we kept all parameters in the models, even if they were not significant. In order to make parameter values for *current diversity* and *natural diversity* comparable, the neighborhoods in the SAR analyses that minimized the AIC for the corresponding *total natural diversity* were used for c*urrent diversity* and *natural diversity of historically extant species*.

## (A) Results

### (B) Diversity gradients

We here focus on the patterns for all terrestrial species (n = 4465) and terrestrial megafauna (n = 330). Three other datasets (large terrestrial species ≥10 kg (n=570) with results similar to those for megafauna; non-marine species (n= 5635), and all species (n=5747) with results overall similar to all terrestrial species) are reported in the appendix (Fig S1-S10). Overall the differences between current and natural diversities were substantially larger for megafauna than for all terrestrial species (Fig. 1). The changes in species, phylogenetic and functional diversity were overall similar although there were some differences which we will discuss later.

**Figure 1.**
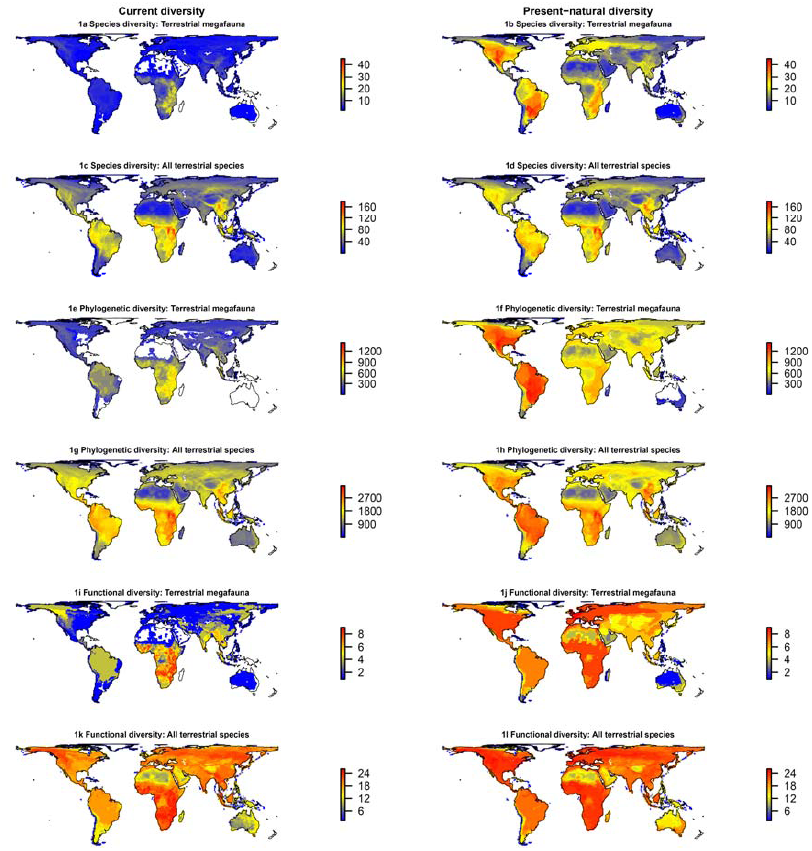
Current diversity and natural species, phylogenetic, and functional diversities for all terrestrial mammal species and for terrestrial mammal megafauna (body size > 10 kg). Colors are standardized horizontally so the same values in are given the same color in all panels.

### (B) Geographic variation in diversity decifits

Geographic patterns in the difference between current and natural diversity (hereafter for simplicity referred to as deficits) in species, phylogenetic and functional diversity exhibit similar geographic patterns (Fig. 2). The largest deficits in megafauna diversity occur on islands (Madagascar, Caribbean, Oceania) and the island-like continent Australia (Fig. 2). Strong deficits are also found in the Americas and Greater Sahara, whereas deficits are only minor in Africa and tropical Asia and intermediate in the remaining regions. The temporal patterns of the losses behind these deficits are radically different. Some regions, such as Australia, New Guinea and the Caribbean islands, and the New World, have had almost exclusively pre-historic losses, others have had mainly historic losses, such as Greater Sahara and Africa, whereas still other regions - most noticeably Europe – have had both large historic and pre-historic losses. In all regions, the relative deficits for all terrestrial species are substantially smaller than the megafauna losses, but the difference between the two varied among regions.

**Figure 2.**
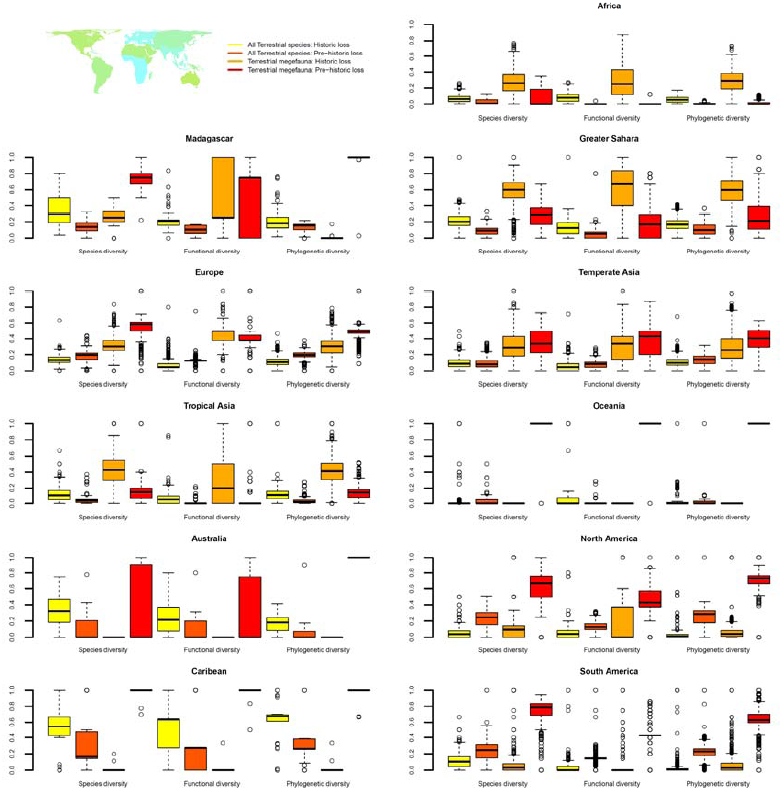
Estimated deficits in natural species, phylogenetic, and functional diversity for all terrestrial mammal species and terrestrial mammal megafauna species. The deficits are divided into those resulting from historic losses (the difference between current diversity and the present natural diversity of all species accepted by IUCN) and those resulting from pre-historic losses (the difference between the natural diversity for all species and for species accepted by IUCN). The thick middle line and box represent the median and first to third quartiles, respectively, and whiskers extend to the furthest datapoint that is no more than 1.5 times the interquartile range away from the median.

### (B) Biases in the inference in diversity drivers

The substantial and geographically variable anthropogenic diversity deficits could have large effects on our ability to understand macroscale diversity patterns. In order to assess the magnitude of this problem, we compared the results of parallel standard macroecological analyses of current and natural diversity patterns. A striking result was that the explanatory power (pseudo-R^2^) was consistently lower for current diversity than for natural diversity for all studied mammal groups. In addition, the models for natural diversity based only on historically extant species consistently had intermediate pseudo-R^2^ values between the two (Tables S1-S3). The decreases in pseudo-R^2^ were especially large for terrestrial megafauna; the pseudo-R^2^ values for all three diversity measures were approximately 0.2 lower for current diversity than for natural diversity. The higher explanatory power for the natural diversity is especially noteworthy given that these are known with less certainty than the current diversity, and these uncertainties would be expected to reduce the pseudo-R^2^ values.

Two other consistent changes were seen in analyses of current vs. natural diversity. In all analyses, the normalized difference vegetation index (NDVI), an indicator of vegetation productivity (Wang *et al*., 2004), was the strongest predictor of diversity, with higher diversity in areas with higher NDVI, but its predictive power was always lower for current diversity than for natural diversity. Elevation range was a weaker, but consistent, predictor (with higher diversity with higher elevation range), but with a consistently stronger effect on current diversity relative to natural diversity. The changes in pseudo-R^2^, NDVI, and elevation range were strikingly regular, with a larger change in pseudo-R^2^ also corresponding to a larger change in the effect sizes of NDVI and elevation range (Fig. 3).

**Figure 3.**
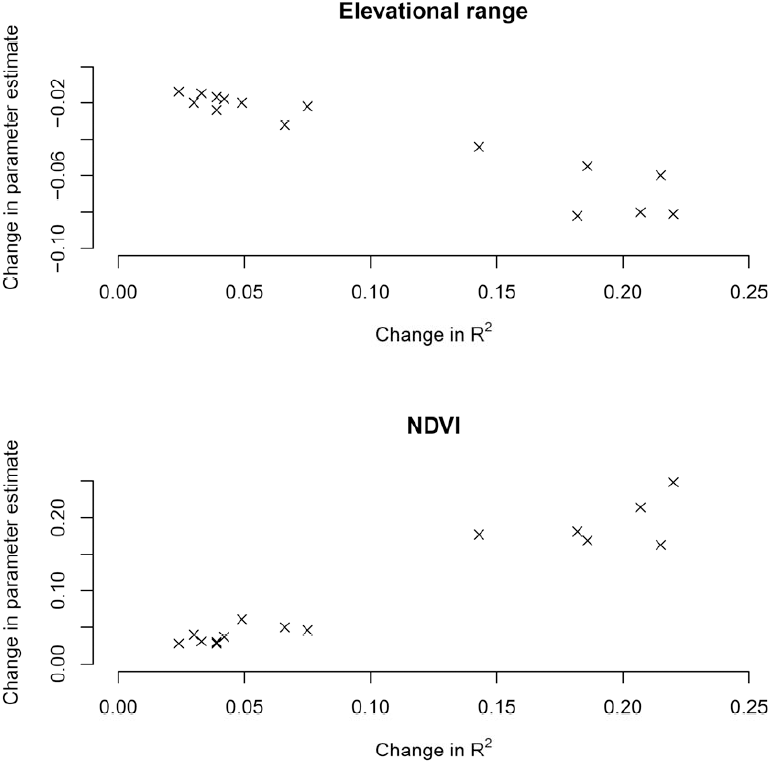
Relationship between the difference in pseudo-R^2^ for the models of natural and current mammal diversity and the corresponding difference between the standardized estimates for the effect size of NDVI or elevation range. The 15 circles represent the difference for each combination of one of the three diversities (species diversity, phylogenetic diversity, functional diversity) and one of the five datasets (all species, non-marine species, all terrestrial species, large terrestrial species, and terrestrial megafauna).

## (A) Discussion

### (B) Diversity gradients

For terrestrial megafauna, the pattern in natural species diversity is radically different from the current pattern (Fig. 1A and B). Current species diversity exhibits a well-known peak in Sub-Saharan Africa, whereas Africa’ s natural species diversity is similar to other continents, as suggested previously (Owen-Smith, 2013). For natural diversity, the highest values are observed in the southern Rocky Mountains and Mexico and in northern Argentina, whereas most of the Americas and large parts of Eurasia have diversities similar to the most diverse areas in sub-Saharan Africa (Fig. 1A and B). Differences between current and natural diversities for all terrestrial species (i.e., irrespective of body size) are smaller than for megafauna, with the largest changes occurring on islands (including the island-like continent Australia) and some temperate areas in Europe and North America (Fig. 1C and D). The patterns in phylogenetic diversity are similar to the patterns in species diversity (Fig. 1E-H versus Fig. 1A-D). The major difference is that natural phylogenetic diversity is elevated in the Americas relative to Africa and Southeast Asia, reflecting greater diversity at deep phylogenetic levels in South America, as seen by tabulating the number of mammalian orders containing terrestrial megafauna.

The natural and current megafauna in Africa belong to six orders, whereas the natural megafauna diversity in South America belongs to nine orders, only five of which have extant megafauna species in South America. The high natural phylogenetic diversity of the New World can likely be seen as a consequence of the former isolation of South America, followed by the effects of the Great American Biotic Interchange (GABI) (Simpson, 1980). Even though many of the formerly endemic South American mammalian groups went extinct due to competition with invading Northern Hemisphere clades, a number of groups survived until the Late Pleistocene or early Holocene (Sandom *et al*., 2014). However, the species diversity within many of these clades was low during the Late Pleistocene, with only a few species, even though they were formerly diverse clades (Billet, 2011), creating a pattern of long branches separating species and high phylogenetic diversity. The GABI is expected to influence the phylogenetic diversity, but it could potentially also influence species diversity. As already suggested by Darwin (1859), related species are often thought to compete more with each other, and two areas of equal productivity may potentially support more species if they are distantly related rather than closely related. The evidence for this is limited (Cahill *et al*., 2008), but a functional coupling between the high megafauna ordinal diversity in the Americas and the region’ s very high species diversity in the absence of human-driven extinctions and extirpations is still possible.

The most striking difference between the patterns in functional diversity (Fig. 1I-L) and the other patterns is that the differences in functional diversity between current and natural diversity are clearly visible in all areas for both all terrestrial species and megafauna, as opposed to being much more evident for megafauna. This is at least partly a logical consequence of the highly size-selective nature of the extinctions and range contractions affecting our size-based metric for functional diversity. Though we mainly focus on patterns that are different between natural and current patterns, the functional diversity patterns also highlight the constancy of some patterns. One such constant pattern is the relative steepness of the gradient between temperate and tropical regions of functional and species diversity. For both current and natural diversity, the gradient is substantially less steep in functional diversity than in the other diversity measures, which corresponds to studies from other taxa (Mouillot *et al*., 2014).

### (B) Geographic variation in diversity decifits

The regional differences in faunal deficits are consistent with higher sensitivity of island regions and open areas to human pressures compared to continental forest regions. Only some of the island regions (Caribbean, Madagascar, and Australia) have large deficits among all terrestrial species, with Oceania having a much smaller deficit, similar to that of tropical Asia. The cause for the low deficits in Oceania is unknown, but it could be related to a higher survival in closed forest environments (Johnson, 2002) due to lower accessibility to humans. The large deficits in the Greater Sahara (Fig. 2) and the desert regions of Australia (Figs. S6-S10) despite low human footprint in these regions (Wildlife Conservation Society, 2015) point to a higher sensitivity of arid faunas to human-driven extirpation either directly via hunting or indirectly via higher sensitivity of arid vegetation to anthropogenic degradation. There is evidence for high pre-historic or historic hunting pressure in certain arid regions, such as the Middle East (Bar-Oz *et al*., 2011). Further, a number of species formerly inhabiting these regions used were at their environmental extremes here, and such marginal populations could potentially be more sensitive to increased human pressures.

Anthropogenic diversity losses in an area within a given time period may depend on losses during preceding periods. Africa experienced a rather large Early Pleistocene extinction, which was potentially caused by early *Homo* species (Werdelin & Lewis, 2013), and this has been suggested to be a contributing factor to the low loss of megafauna in Africa in the Late Pleistocene (Short & Smith, 1994). Similarly, the relatively low historic losses in North and South America may reflect that the massive pre-historic loss already had removed most sensitive species. Conversely, the lower pre-historic loss in South-East Asia and potential survival of relatively sensitive species could explain its large historic losses and. higher fraction endangered species compared to America (Sodhi *et al*., 2010).

The patterns in phylogenetic diversity deficits are similar to those for species diversity, although generally smaller (Fig. 2). This is most striking in Australia, which appears to have suffered substantially lower phylogenetic losses relative to its species losses. A potential reason may be that marsupials appear to have a higher evolutionary ecological plasticity than placental mammals, so that phylogenetic clades are less ecologically specialized and thus less consistently sensitive to the same pressures. Exemplifying this, the two largest extinct marsupial predators belongs to two different orders (Dasyuromorphia and Diprotodontia), with the latter showing large variation in body size, from <10 g to >1 ton. The cause of this large plasticity is unknown, but it could potentially be a corollary of the more limited scope for ecological specialization in marsupials caused by only having one set of teeth (Werdelin, 1987). Irrespective of the underlying cause, selective removal of species based on ecological characteristics, such as body size, would remove a relatively lower amount of phylogenetic history for marsupials than for placental mammals.

### (B) Biases in the inference in diversity drivers

The findings of this paper suggest that analyses of current patterns may lead to a biased understanding of the drivers of diversity. Importantly, our results suggest that mammal diversity was more strongly linked to vegetation productivity before being reshaped by human activities and that analyses of current diversity patterns underestimate this relationship. This is likely a consequence of the strong correlation between productivity and human population density (Evans & Gaston, 2004). The differences in the importance of elevation range also suggest an anthropogenic bias in the current patterns. All current diversities exhibited significant positive correlations between diversity and elevation ranges, whereas this relationship was not significant for the natural diversity of megafauna and large species diversity. Habitat accessibility for humans may be highly correlated with elevation range as suggested by a strong correlation between elevation slope and remaining tree cover across the globe (Sande & Svenning, 2013). Importantly, steep mountainous terrain has been proposed to have offered some megafauna species a refuge from pre-historic hunting (Johnson, 2002) and to provide protection against human activities today (Gavashelishvili & Lukarevskiy, 2008). Therefore, our results suggest that the positive effect of elevation range on the current diversity of terrestrial megafauna may largely be an anthropogenic artefact rather than a natural phenomenon, as it is not a significant predictor of natural diversity. On the other hand, the still significant effect of elevational range for the natural diversity of all terrestrial species (Tables S1-S3) suggest that the effect on overall diversity represent a natural phenomenon, e.g., reflecting habitat heterogeneity.

The performance of models of diversity drivers is often judged based on their explanatory power (Jetz & Fine, 2012), and changes in R^2^ values may change our understanding of how well we understand diversity patterns. There is still substantial debate over the causes of the overall diversity gradients (Brown, 2014). Part of the reason why no explanation has been universally accepted could be that the models are designed to explain natural diversity, but are applied to current diversity. The observed increases in pseudo-R^2^ values when shifting to natural diversity suggest that we are actually better able to explain diversity gradients than we thought, at least for mammals, if we remove the human-induced biases.

We have only focused on overall diversity patterns rather than clade- or region-specific patterns, but there is no reason to assume that the overall patterns are especially sensitive to human impact. One could even assume that the patterns we investigated in this paper may be some of the geographic patterns least influenced by anthropogenic modifications. Many studies have documented a strong human impact on smaller-scale distribution and diversity patterns (Laliberte & Ripple, 2004). Therefore, we suggest that researchers working on macroecological or macroevolutionary analyses of natural ecological and evolutionary drivers of diversity should focus on natural rather than current distributions whenever possible (maps of all distributions are available as appendix 2-7, but the data can also be downloaded in a readily useable format at http://bios.au.dk/om-instituttet/organisation/oekoinformatik-biodiversitet/data/ (DATAWILL BE UPLOADED UPON ACCEPTANCE).

More fundamentally, the results of the present study illustrate that we now live in the Anthropocene (Crutzen, 2002), a human-dominated epoch in which few biological patterns and processes are not substantially modified by humans (Helmus *et al*., 2014; Dirzo *et al*., 2014). Therefore, it is important to integrate the potential effects of humans into any type of analysis, including ones often thought of as being little influenced by humans, such as the diversity gradients discussed here. The estimated natural distributions for all late-Quaternary mammals will also be highly useful for applied conservation projects and studies, enabling managers and researchers to use present natural diversity as a baseline, e.g., for selecting species for rewilding projects or reintroductions (Donlan *et al*., 2005; Hayward 2009).

## Supplementary Materials

Supplementary Materials and Methods

Supplementary Figures S1-S11

Supplementary Tables S1-S3

Supplementary Data 1-8

## Acknowledgments

JCS was supported by the European Research Council (ERC-2012-StG-310886-HISTFUNC). SF was supported by the Danish Natural Science Research Council (#11-115750). SF estimated the natural distributions of all species with input from JCS. SF performed the analyses. SF and JCS wrote the paper.

## Data accessibility

The data reported in this paper are provided in the electronic supplementary material as pdf files and in a readily useable format from http://bios.au.dk/om-instituttet/organisation/oekoinformatik-biodiversitet/data/ (DATAWILL BE UPLOADED UPON ACCEPTANCE).

## Biosketches

**Søren Faurby** is a postdoc at Aarhus University. He is an evolutionary biologist interested in the development, maintenance, and consequences of geographic variation within and between species, and he has investigated these subjects in a wide variety of taxa.

**Jens-Christian Svenning** is a professor at Aarhus University. He is a broadly based ecologist, with core research interests including community and vegetation ecology, macroecology, biogeography, and physical geography. His work ranges from addressing basic ecological and evolutionary questions to investigating applied ecology, conservation biology, and global change.

## REFERENCES

Álvarez-Lao, D.J. & García, N. (2010) Chronological distribution of Pleistocene cold-adapted large mammal faunas in the Iberian Peninsula. Quaternary International, 212,120&128.

Bar-Oz, G., Zeder, M. & Hole, F. (2011) Role of mass-kill hunting strategies in the extirpation of Persian gazelle (*Gazella subgutturosa*) in the northern Levant. Proceedings of the National Academy of Sciences, 108, 7345&7350.

Barnosky, A.D., Koch, P.L., Feranec, R.S., Wing, S.L. & Shabel, A.B. (2004) Assessing the causes of Late Pleistocene extinctions on the continents. Science, 306, 70&75.

Barnosky, A.D., et al (2011) Has the Earth’s sixth mass extinction already arrived? Nature, 471, 51&57.

Biknevicus, A.R., McFarlane, D.A. & MacPhee, R.D.E. (1993) Body size in *Amblyrhiza inundata* (Rodentia: Caviomorpha), an extinct megafaunal rodent from the Anguilla Bank, West Indies: estimates and implications. American Museum Novitates, 3079, 1&25.

Billet, G. (2011) Phylogeny of the Notoungulata (Mammalia) based on cranial and dental characters. Journal of Systematic Palaeontology, 9, 481&497.

Blackburn, T.M. & Gaston, K.J. (1998) Methodological issues in macroecology. American Naturalist, 151, 68&83.

Bover, P.P. & Alcover, J.A. (2008) Extinction of the autochthonous small mammals of Mallorca (Gymnesic Islands, Western Mediterranean) and its ecological consequences. Journal of Biogeography, 35, 1112&1122.

Brown, J.M. (2014) Why are there so many species in the tropics? Journal of Biogeography, 41, 8&22.

Cahill, J.F., Kembela, S.W., Lamba, E.G. & Keddy, P.A. (2008) Does phylogenetic relatedness influence the strength of competition among vascular plants? Perspectives in Plant Ecology, Evolution and Systematics, 10, 41&50.

Crutzen, P.J. (2002) Geology of mankind. Nature, 415, 23&23.

Darwin, C. (1859) The Origin of Species. John Murray, London.

De Thoisy, B., Richard-Hansen, C., Goguillon, B., Joubert, P., Obstancias, J., Winterton, P. & Brosse, S. (2010) Rapid evaluation of threats to biodiversity: human fotprint score and large vertebrate species responses in French Guiana. Biodiversity Conservation, 19,1567&1584.

Devictor, V., Mouillot, D., Meynard, C., Jiguet, F., Thuiller, W. & Mouquet, N. (2010) Spatial mismatch and congruence between taxonomic, phylogenetic and functional diversity: the need for integrative conservation strategies in a changing world. Ecology Letters, 13, 1030&1040.

Dirzo, R., et al(2014) Defaunation in the Anthropocene. Science, 345, 401&406.

Donlan, J., et al. (2005) Re-wilding North America. Nature, 436, 913&914.

Duncan, R.P., Boyer, A.G. & Blackburn, T.M. (2013) Magnitude and variation of prehistoric bird extinctions in the Pacific. Proceedings of the National Academy of Sciences, 110, 6436&6441.

Evans, K.L. & Gaston, K.J. 2004 People, energy and avian species richness. Global Ecology and Biogeography, 14, 187&196.

Faurby, S. & Svenning, J.C. (2015) A species-level phylogeny of all extant and late Quaternary extinct mammals using a novel heuristic-hierarchical bayesian approach. Molecular Phylogenetics and Evolution, 84, 14&26.

Gavashelishvili, A. & Lukarevskiy, V. (2008) Modelling the habitat requirements of leopard *Panthera pardus* in west and central Asia. Journal of Applied Ecology, 45, 579&588.

Hayward, M.W. (2009) Conservation management for the past, present and future. Biodiversity and Conservation, 18,765&775.

Helmus, M.R., Mahler, D.L. & Losos, J.B. (2014) Island biogeography of the Anthropocene. Nature 513, 543&546.

Hijmans, R.J., Cameron, S.E., Parra, J.L., Jones, P.G., Jarvis, A. (2005) Very high resolution interpolated climate surfaces for global land areas. International Journal of Climatology, 25, 1965&197.

Huang, S., Stephens, P.R. & Gittleman, J.L. (2012) Traits, trees and taxa: global dimensions of biodiversity in mammals. Proceedings of the Royal Society B, 279, 4997&5003.

IPCC (2013) *Climate Change 2013: The Physical Science Basis. Contribution of Working Group I to the* Fifth Assessment Report of the Intergovernmental Panel on Climate Change (ed. by Stocker, T.F., et al.). Cambridge University Press, Cambridge.

Jetz, W. & Fine, P.V.A. (2012) Global gradients in vertebrate diversity predicted by historical area-productivity dynamics and contemporary environment. PLoS Biology, 10, e1001292.

Johnson, C.N. (2002) Determinants of loss of mammal species during the Late Quaternary ‘megafauna’ extinctions: life history and ecology, but not body size. Proceedings of the Royal Society of London B, 269, 2221&2227.

Kissling, W.D. & Carl, G. (2008) Spatial autocorrelation and the selection of simultaneous autoregressive models. Global Ecology and Biogeography, 17, 59&71.

Koch, P.L. & Barnosky, A.D. (2006) Late Quaternary extinctions: State of the debate. Annual Reviews of Ecology Evolution and Systematics, 37, 215&250.

Laliberte, A.S. & Ripple, W.J. (2004) Range contractions of North American carnivores and ungulates. BioScience, 154, 123&138.

Mazel, F., et al. 2014 Multifaceted diversity–area relationships reveal global hotspots of mammalian species, trait and lineage diversity. Global Ecology and Biogeography, 23, 836&847.

Mouillot, D., et al. (2014) Functional over-redundancy and high functional vulnerability in global fish faunas on tropical reefs. Proceedings of the National Academy of Sciences, 111, 13757&13762.

Nowak, R.M. (1999) Walker’s Mammals of the World. The Johns Hopkins University Press, Baltimore.

Olson, D.M. (2001) Terrestrial ecoregions of the world: A new map of life on Earth. BioScience, 51, 933&938.

Owen-Smith, N. (2013) Contrasts in the large herbivore faunas of the southern continents in the late Pleistocene and the ecological implications for human origins. J. Biogeogr. 40, 1215&1224.

Pennisi, E. (2005) What determines species diversity? Science, 309, 90.

Peterken, G.F. (1977) Habitat conservation priorities in British and European woodlands. Biodiversity Conservation, 11, 223&236.

Ripple, W.J., et al. (2014) Status and ecological effects of the world’s largest carnivores. Science, 343, 1241484 (2014).

Safi, K., Cianciaruso, M.V., Loyola, R.D., Brito, D., Armour-Marshall, K. & Diniz-Filho, J.A.F. (2011) Understanding global patterns of mammalian functional and phylogenetic diversity. Philosophical Transactions of the Royal Society B, 366, 2536&2544.

Sandel, B. & Svenning, J.C. (2013) Human impacts drive a global topographic signature in tree cover. Nature Communications 4, 2474.

Sandom, C., Dalby, L., Fløjgaard, C., Kissling, W.D., Lenoir, J., Sandel, B., Trøjelsgaard, K., Ejrnæs, R. & Svenning, J.C. (2013) Mammal predator and prey species richness are strongly linked at macroscale. Ecology, 94, 1112&1122.

Sandom, C., Faurby, S., Sandel, B. & Svenning, J.C. (2014) Global late Quaternary megafauna extinctions linked to humans, not climate change. Proceedings of the Royal Society B,. 281, 20133254.

Schipper, J. et al. (2008) The status of the world’s land and marine mammals: Diversity, threat, and knowledge. Science, 322, 225&230.

Short, J., Smith, A. (1994) Mammal decline and recovery in Australia. Journal of Mammalogy, 75, 288&297.

Simpson, G.G. (1980) *Splendid isolation: the curious history of South American mammals*. Yale University Press.

Sodhi, N.S., Posa, M.R.C., Lee, T.M., Bickford, D., Koh, L.P. & Brook, B.W. (2010) The state and Conservation of Southeast Asian biodiversity. Biodiversity Conservation, 19, 317&218.

Sommer, R.S., Benecke, N. (2006) Late Pleistocene and Holocene development of the felid fauna (Felidae) of Europe: a review. Journal of Zoology. 269, 7&19.

Spear, D. & Chown, S.L. (2008) Taxonomic homogenization in ungulates: patterns and mechanisms at local and global scales. Journal of Biogeography, 35, 1962&1975.

Steadman, D.W., Martin, P.S., MacPhee, R.D.E., Jull, A.J.T., McDonald, H.G., Woods, C.A., Iturralde-Vinent, M. & Hodgins, G.W.L. (2005) Asynchronous extinction of late Quaternary sloths on continents and islands. Proceedings of the National Academy of Sciences, 102, 11763&11768.

Tucker, C.J., Pinzon, J.E., Brown, M.E., Slayback, D.A., Pak, E.W., Mahoney, R., Vermote, E.F. & El Saleous, N. (2005) An Extended AVHRR 8-km NDVI Data Set Compatible with MODIS and SPOT Vegetation NDVI Data. International Journal Remote Sensing, 26, 4485&4498.

Turvey, S.T. & Fritz, S.A. (2011) The ghosts of mammals past: biological and geographical patterns of global mammalian extinction across the Holocene. Philosophical Transactions of the Royal Society B, 366, 2564&2576.

UCLA: Statistical Consulting Group. (2014) FAQ: What are pseudo R-squareds? (Available at http://www.ats.ucla.edu/stat/mult_pkg/faq/general/Psuedo_RSquareds.htm). (Accessed: 12th February 2015).

Wang, J., Rich, P.M., Price, K.P. & Kettle, W.D. (2004) Relations between NDVI and tree productivity in the central Great Plains. International Journal of Remote Sensing, 25, 3127&3138.

Werdelin, L. (1987) Jaw geometry and molar morphology in marsupial carnivores: analysis of a constraint and its macroevolutionary consequences. Paleobiology, 13, 342&350.

Werdelin, L & Lewis, M.E. (2013) Temporal change in functional richness and evenness in the eastern African Plio-Pleistocene carnivoran guild. PLoS ONE, 8, e57944.

Wildlife Conservation Society - WCS, and Center for International Earth Science Information Network - CIESIN - Columbia University. (2015) Last of the Wild Project, Version 2, 2005 (LWP-2): Global Human Footprint Dataset (Geographic). (Available at http://dx.doi.org/10.7927/H4M61H5F). (Accessed: 12th February 2015).

Yeakel, J.D. (2014) Collapse of an ecological network in Ancient Egypt. Proc. Natl. Acad. Sci. U.S.A. 111, 14472&14477.

